# An accessible digital imaging workflow for multiplexed quantitative analysis of adult eye phenotypes in *Drosophila melanogaster*

**DOI:** 10.1101/2024.01.26.577286

**Authors:** Heidi J. J. Pipkin, Hunter L. Lindsay, Adam T. Smiley, Jack D. Jurmu, Andrew M Arsham

## Abstract

The compound eye of *Drosophila melanogaster* has long been a model for studying genetics, development, neurodegeneration, and heterochromatin. Imaging and morphometry of adult *Drosophila* and other insects is hampered by the low throughput, narrow focal plane, and small image sensors typical of stereomicroscope cameras. When data collection is distributed among many individuals or extended time periods, these limitations are compounded by inter-operator variability in lighting, sample positioning, focus, and post-acquisition processing. To address these limitations we developed a method for multiplexed quantitative analysis of adult *Drosophila melanogaster* phenotypes. Efficient data collection and analysis of up to 60 adult flies in a single image with standardized conditions eliminates inter-operator variability and enables precise quantitative comparison of morphology. Semi-automated data analysis using ImageJ and R reduces image manipulations, facilitates reproducibility, and supports emerging automated segmentation methods, as well as a wide range of graphical and statistical tools. These methods also serve as a low-cost hands-on introduction to imaging, data visualization, and statistical analysis for students and trainees.

## Introduction

The foundational studies of plant and animal genetics relied on visible morphological traits to reveal the function and inheritance of genes – in peas, flies, maize, and mice, traits defined by size, shape, color, and texture illuminated for the first time the existence and behavior of genes, chromosomes, transposons and more. Tools to measure the molecular processes underlying these traits are now abundant and powerful, but the importance of physical traits persists, as phenotypes *per se* and as indirect reporters of genetic interactions. In contrast to the explosive pace of change in molecular techniques, whole-animal imaging has changed slowly and is hampered by a lack of scale: most microscope image sensors have small fields of view and narrow focal planes that photograph a small number of mostly out-of-focus animals. Small working distances on compound microscopes and inconsistent lighting on stereomicroscopes are additional barriers.

Focus stacking or z-stacking can generate high quality images of 3 dimensional samples for entomology collections (Droege and Gutierrez 2024), and for cataloging of phenotypes (Holtzman and Kaufman 2013). Many Drosophilists’ first act as PI is to hang the *Learning to Fly* poster (Childress et al. 2005) in their fly rooms as a visual reference of common phenotypes, but for most labs creating such images of specific phenotypes of interest is out of reach.

Since Thomas Hunt Morgan isolated the first white-eyed mutant of *Drosophila melanogaster* (Morgan 1910) and Hermann J. Muller generated heterochromatin-silenced chromosomal inversions (Muller 1930), studies of the fly eye have made foundational contributions to the understanding of gene expression and development. Among these is a century of work to understand heterochromatin and its role in gene regulation (Elgin and Reuter 2013). Fly eye phenotypes are also powerful tools for studying neurodegeneration (McGurk et al. 2015) and for identifying causal mutations in (and possible treatments of) human disease (Dalton et al. 2022; Manivannan et al. 2022). Spectrophotometric measurement of eye pigment from homogenized flies is quantitative (Huisinga et al. 2016) but collapses inter- and intra-individual variation into a single value. Photographic measurement of eye color is quantitative and captures variation in patterns and levels (Diez-Hermano et al. 2015; Iyer et al. 2016; Swenson et al. 2016; Kelsey and Clark 2017; Diez-Hermano et al. 2020) but to date has had limited throughput and resolution.

Here we describe a cost-effective method for multiplexed quantitative analysis of up to 60 adult *Drosophila* in a single image using a full-frame digital camera and macro lens on a motorized rail. Focus stacking combines the sharpest pixels of each photo into a single composite image. Semi-automated data extraction and analysis using ImageJ and R facilitate code sharing and reduce intermediary data products. Inter-operator variability among early career researchers with a wide range of experience was less than 6%, and often much lower, demonstrating the suitability and robustness of these techniques for a variety of research and educational settings.

## Materials and Methods

### Stocks used

90 Stocks used
| Name | Genotype | Source |
| --- | --- | --- |
| Dorsal eye | y[1] w[*]; wg[Sp-1]/CyO;<br>P{w[+mW.hs]=GawB}mirr[DE]/TM3, Sb[1] | BDSC 29650 (Morrison and Halder 2010) |
| lacO 010 | yw; P{y+ lacO hsp70-white} | This study; P element mobilization of 1198-lacO from the Elgin lab |
| lacO 025 | yw; P{y+ lacO hsp70-white} | This study; P element mobilization of 1198-lacO from the Elgin lab |
| wm4 | y ln(1) w<sup>m4</sup> | Kind gift of Elgin lab (Muller 1930) |
| wt 1 | Maple Grove Wild-Type | This study; isolated in Heidi J.J. Pipkin's kitchen, Maple Grove, MN |
| wt 2 | RAL365 | The Drosophila Genetic Reference Panel (Mackay et al. 2012) |
| eya composite-GAL4 w+ | yw; ;eya composite-GAL4 w+ | Kind gift of Justin Kumar (Weasner et al. 2016) |
| eya composite-GAL4 w- | yw; ;eya composite-GAL4 w- | This study; *white* reporter minigene mutated with CRISPR |
| mini-white | yw*; ; eyGAL4 | Kind gift of Justin Kumar (Weasner et al. 2016) |
| White eraser | w[*]; P{y[+t7.7] GFP[3xP3.cUa]=white-<br>eraser}attP2 | BDSC 90371 (Liu et al. 2020) |
| HP1 RNAi | y[1] sc[*] v[1] sev[21]; ;P{y[+t7.7]<br>v[+t1.8]=TRiP.HMS00278}attP2 | BDSC 33400 (Zirin et al. 2020) |
| CG17359 RNAi | y[1] v[1]; P{y[+t7.7]<br>v[+t1.8]=TRiP.JF02340}attP2 | BDSC 26776 (Zirin et al. 2020) |
| CG17361 RNAi | y[1] sc[*] v[1] sev[21]; P{y[+t7.7]<br>v[+t1.8]=TRiP.HMC04903}attP40 | BDSC 57714 (Zirin et al. 2020) |

#### Fly husbandry and crosses

Where indicated, fly stocks were obtained from the Bloomington Drosophila Stock Center (BDSC, RRID:SCR_006457). Most GAL4 stocks use a dominant *white* transgene as a positive selection marker for the presence of GAL4, preventing measurement of *white* reporter gene expression. We used “white eraser” flies expressing Cas9 and guide RNAs targeting *white* (Liu et al. 2020) to eliminate *mini-white* expression with CRISPR. Female “white eraser” flies were crossed with males carrying GAL4 driven by a composite enhancer constructed from regulatory elements of the *eyes absent (eya)* gene (Weasner et al. 2016) on the third chromosome. Male flies with *eya composite-GAL4* (hereafter referred to as *eya-GAL4*) and *P*{*white eraser*} were crossed to third chromosome balancer stocks, and individual *white* males with *eya-GAL4* but without the fluorescent markers indicating the presence of *P*{*white eraser*} were used to establish new white-eyed *eya-GAL4* stocks. These stocks were then crossed to *yellow* flies carrying the X-ray-induced *w*^*m4*^ inversion (Muller 1930) to generate the driver-reporter stock *y w*^*m4*^; *;eya-GAL4*^*w-*^ (hereafter abbreviated *w*^*m4*^; *eya-GAL4*).

All crosses were incubated on standard Bloomington media (Nutri-Fly BF, Genesee Scientific) at 25°C and at least 60% relative humidity. For generation of eye color variation, 2-4 males of each eye pigment stock were crossed to 6-10 unmated *yw* females in 3 independent vials. For RNAi knockdown experiments, unmated *w*^*m4*^; *eya-GAL4* females were crossed with males expressing RNAi against the gene of interest (Zirin et al. 2020). For all crosses, 2-4 day old adult progeny were collected and quickly frozen at -20°C for later analysis.

We used FlyBase release FB2024_03 to obtain information on gene structure and expression ((Öztürk-Çolak et al. 2024) RRID:SCR_006549). Images, data, code, and design files for this study (Arsham 2024) are available on FigShare (RRID:SCR_004328).

#### Image acquisition

Frozen flies were thawed and arranged on a custom-designed grooved 3D-printed sample tray with a color control to ensure consistency between images (Matte ColorGauge Pico, #87-414, Edmund Optics). The sample tray was then placed in a 3D printed 360° lighting system with 120 white LEDs (color temperature 6000K) diffused by a strip of translucent white mylar on an adjustable XY stage centered under the image sensor (Figure 1). A parts list and all design files for lighting and mounting can be found at https://www.thingiverse.com/thing:4688444 and https://www.thingiverse.com/thing:4596690 respectively.

**Figure 1.**
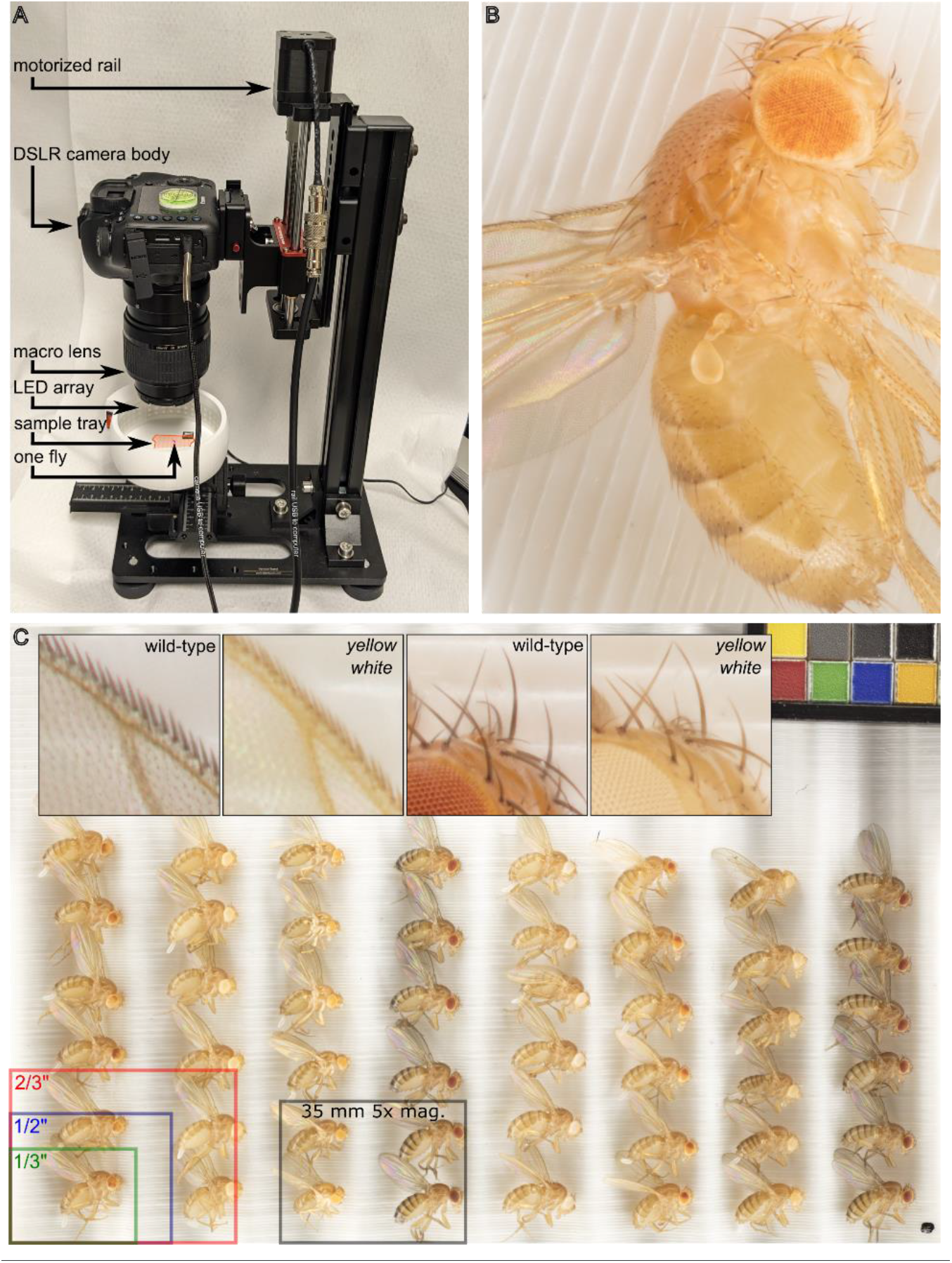
A digital photography workflow for multiplexed analysis of Drosophila melanogaster phenotypes. (**A**) Canon EOS 5Ds camera body and macro lens mounted to motorized rail and mounted on a vertical stand. Camera and rail are independently connected to and remote-controlled from a Windows computer by USB cables. Flies are positioned on a 3D printed grooved sample tray (highlighted in orange) and illuminated by 360° LED lighting system on a manually adjustable XY stage. A single fly, highlighted in pink, is pictured to illustrate scale. (**B**) Focus-stacked 5x magnification image of Drosophila melanogaster with PEV phenotype. (**C**) Full frame image of 48 adult animals of various genotypes as well as a color-checker (top right) to ensure consistency. Top inset, from left to right, 5x magnification images of wild-type and yellow mutant wing edge, and wild-type and yellow white ocellus and head bristles. Bottom overlay, areas of common microscope camera image sensor formats and of our system at 5x magnification.

We used a Canon MP-E 65 mm f/2.8 1-5x Macro lens attached to a Canon EOS 5Ds camera with a 50 megapixel 36 × 24 mm CMOS sensor, pixel pitch of 4.13 µm and pixel area of 17.06 µm^2^. The camera and a vertical rail (WeMacro 100 mm rail #WM001 and vertical stand #WVH01) were controlled from a computer running Windows 10 and Helicon Remote version 3.9.11. Images were acquired in RAW format with exposure of 1/25 s, f/2.8, and either ISO 100 (for 1x magnification) or ISO 500 (to correct for the reduction in light reaching the sensor at 5x magnification). Rail travel from top to bottom is 2750 µm made up of 55 steps at 50 µm each. The 56-image stack was automatically exported from Helicon Remote to Helicon Focus version 7.7.5 and a composite TIF image combining the sharpest areas of each individual image was generated using the “C, smoothing 4” setting. All original quantitated images described here are publicly available (Arsham 2024).

### Data analysis

Individual users defined a region of interest (ROI) for every eye in an image using the elliptical ROI tool in the FIJI distribution of ImageJ (version 1.53). The BAR plugin (version 1.51 (Ferreira et al. 2017)) was used to apply a standardized colon-delimited naming convention to all ROIs specifying sex, genotype, replicate number, user, and other key experimental variables. A custom ImageJ macro converted the image to RGB, inverted the colors so that higher pigment levels (darker red eyes) correspond to higher RGB values, and converted to 8-bit grayscale using ImageJ’s built-in weighted grayscale conversion:

~~~
row = 0; //resets results row
      roiCount = roiManager(“count”);
      for (i=0; i<roiCount; i++) {
            run(“RGB Color”);
            run(“Conversions…”, “scale weighted”);
            run(“8-bit”);
            roiManager(“select”, i);{ //start loop
                 Roi.getBounds(rx, ry, width, height);
                       for(y=ry; y<ry+height; y++) {
                 for(x=rx; x<rx+width; x++) {
                 if(Roi.contains(x, y)==1) {
                       setResult(“ROI”, row, Roi.getName);
                       setResult(“pixel”, row, getPixel(x, y));
                 row++;
     }
    }
   }
 }
}
~~~

Each pixel has a single inverted grayscale color value between 0 (white) and 255 (black) that correlates to eye pigmentation. The ImageJ macro saves the grayscale value of each pixel from each segmented eye into a single CSV file that is saved alongside the original (still unmodified) image file. CSV files are imported into R Studio, and data from multiple images are concatenated into a single data frame for all conditions and replicates. The colon-delimited ROI identifiers contain the experimental conditions for each pixel and are separated into factors so that any pixel can be grouped by sex, genotype, replicate, user, etc. All code and data described here are publicly available (Arsham 2024).

## Results and Discussion

### Principles of computational biology

Our goal was to develop a rigorous and affordable method for multiplexed quantitative analysis of adult *Drosophila melanogaster* phenotypes suitable for a wide range of environments from research-intensive labs to undergraduate classrooms. We focused on minimizing photo manipulations and intermediate data products, and on creating simple, clear paths from data to analysis that can be run by any scientist at any time (Royle 2019). To this end, the only manual step after arranging the flies on the sample tray is to draw an ellipse around each eye in the resulting image in ImageJ. This process, known as image segmentation, converts human visual pattern recognition into a computer-readable list of coordinates. Each step in the process (setting up a cross, collecting and freezing flies, staging and photographing, segmentation, data extraction, and data analysis) can be spread out over time or completed by different people.

The system described here, including all hardware, software, and 3D-printed parts, can be assembled for less than $5,000, substantially less than commercially available “macroscopes.” While proprietary software is used to capture and process the images, all post-acquisition analysis steps use freely available open-source cross-platform software.

### Analysis of color variation in *Drosophila* mutants

Microscope cameras are often optimized for high sensitivity and low noise for light-limited applications like fluorescence or for small flat areas like slide-mounted tissue sections. These trade-offs are poorly suited to applications with abundant illumination and three-dimensional objects like whole insects. Our system optimizes for sensor size and resolution rather than sensitivity and noise. The area of a 35 mm image sensor is 8.4 × 10^8^ microns; Figure 1 shows the approximate size of common microscope image sensors for comparison. A 1:1 or “true macro” image can capture up to 60 flies, with each individual fly comprising about 0.7 megapixels (Figure 1 and Figure 2). At 5x magnification (the maximum optical zoom of our macro lens) phenotypes of individual ommatidia, ocelli, bristles, and wing cells are clearly visible (Figure 1B and 1C inset).

**Figure 2.**
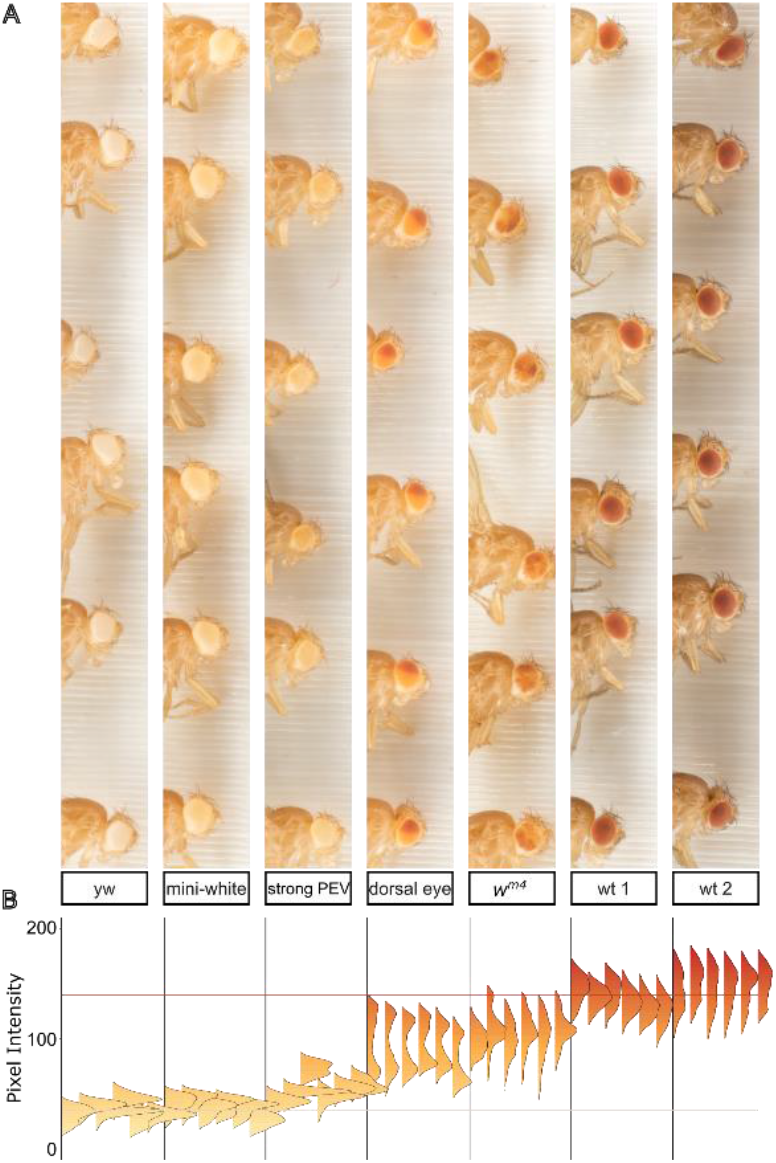
Pigment analysis of eye color mutants. (**A**) A single unmagnified macro image of flies from various stocks was cropped and arranged in order of pigment expression. Individual eyes were segmented in ImageJ and the intensity value of all pixels of each eye are displayed as a vertical histogram (**B**). The beige horizontal line at y = 35 denotes average pixel intensity value for white eyes; the red horizontal line at y = 140 denotes average pixel intensity for wild type.

This system compares favorably in throughput and precision with other computational approaches to eye color (Swenson et al. 2016; Kelsey and Clark 2017), and we tested it on several phenotypically distinct populations of flies. These included flies with no eye pigmentation (*yw)*, a low expression mini-white transgene (mini-white) producing very light yellow eye color, a stock from our lab that expresses a mini-white transgene that is stochastically silenced (strong PEV), the “dorsal eye” stock in which pigmentation is spatially and spectrally bimodal (Morrison and Halder 2010), the *w*^*m4*^ inversion generated by Hermann J. Muller in his studies of the effect of X-rays on inherited phenotypes (Muller 1930), and two wild-type stocks, one isolated locally and one obtained from the *Drosophila melanogaster* Genetic Reference Panel (Mackay et al. 2012), labeled wt 1 and wt 2 respectively. In each case, we crossed males of the indicated stock with *yw* females so that all groups were heterozygous for whichever form of the *white* gene they carried.

Pixel intensity values within or across experiments can be grouped or compared using any experimental variables, and histograms of pigment intensity in individual eyes can reveal bimodal or other non-normal pigment distributions (Figure 2B). To sample and compare multiple populations of flies we generated a single mean pixel intensity value for each eye and a population mean from all the eyes. To visually orient the viewer and provide internal landmarks for comparison, we established average mean eye color values for *white* (pixel intensity = 35) and wild-type (pixel intensity = 140) flies and plotted standard lines on each graph: a beige line to represent *white* eyes and a red line for wild-type.

### Analysis of PEV modification by RNAi knockdown

To apply these approaches to hypothesis-driven experiments we used *in vivo* RNAi (Zirin et al. 2020) to measure the effect of candidate gene knockdown on the *w*^*m4*^ mutant, a well-studied reporter of heterochromatin-mediated gene silencing (Muller 1930) in which the *white* gene is partially silenced by pericentric heterochromatin (Figure 3).

**Figure 3.**
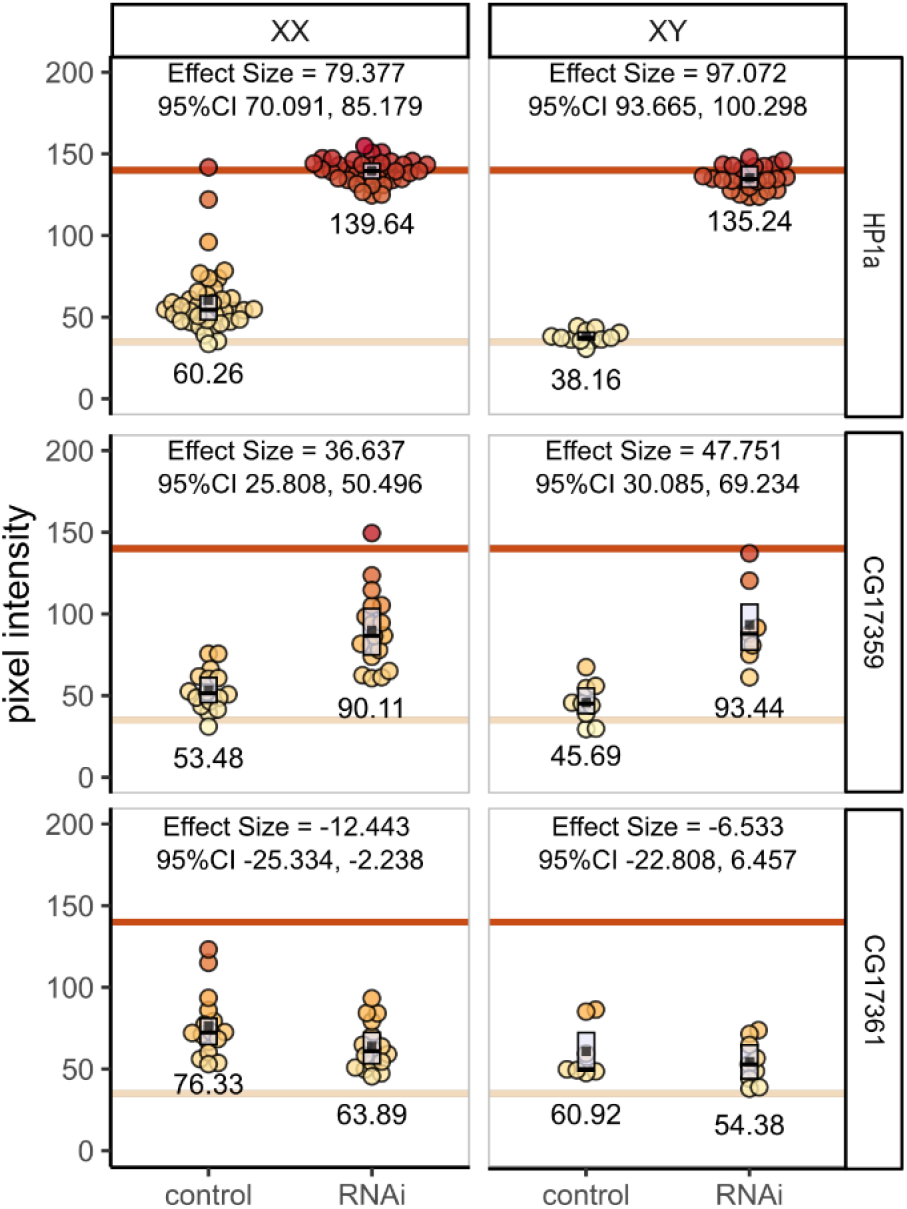
RNAi knockdown of candidate regulators of heterochromatin. Individual fly eyes form at least three independent crosses were photographed and analyzed. Each filled circle is a single eye; group averages are shown numerically and as filled black squares. Boxplots show interquartile range with the median as a black horizontal line. The three RNAi stocks tested are arranged by row; female and male flies are separated by column. Within each panel flies with the w^m4^ mutation and the eyaGAL4 driver but no RNAi are shown on the left; w^m4^, eyaGLA4, and RNAi are shown on the right.

Using eye-specific GAL4 (Weasner et al. 2016) to drive the expression of shRNA, we targeted the essential heterochromatin component HP1a (James and Elgin 1986) and two candidate genes of unknown function by crossing female *w*^*m4*^; *eya-GAL4* flies to male UAS-RNAi stocks (Figure 3). As expected, RNAi knockdown of HP1a abrogated heterochromatin, more than doubling pigmentation in female flies and tripling it in males (mean pigment difference 79.377 [95%CI 70.091, 85.179] and 97.072 [95%CI 93.665, 100.298] in females and males respectively).

The ZAD-ZNF gene family is evolutionarily dynamic (Kasinathan et al. 2020). Several of the 90+ genes in this family regulate heterochromatin (Weiler 2007; Swenson et al. 2016; Baumgartner et al. 2022; Shapiro-Kulnane et al. 2022) but the vast majority are uncharacterized. CG17359 and CG17361 are similar adjacent single-exon genes on chromosome 3L. Both are highly expressed in ovaries (Brown et al. 2014; Leader et al. 2018), and CG17359 was identified as a fast-evolving gene and a strong candidate for heterochromatin regulatory function (Kasinathan et al. 2020). Indeed, when we knocked down CG17359, we observed substantial increases in eye pigment suggesting a disruption of heterochromatin (mean pigment difference 36.637 [95%CI 25.808, 50.496] and 47.751 [95%CI 30.085, 69.234] in females and males respectively) and implicating CG17359 in heterochromatin regulation. In contrast, knockdown of its adjacent paralog CG17361 did not increase pigment levels; we observed a very slight pigment decrease (mean pigment difference -12.443 [95%CI -25.334, - 2.238] and -6.533 [95%CI -22.808, 6.457] for females and males respectively).

### Analysis of inter-operator variability

To assess inter-operator reproducibility of image segmentation and to test the suitability of these methods for early career scientists including undergraduate students, a photo containing flies from each genotype (such as that shown in Figure 2A) was given to 5 undergraduate students ranging in lab experience from a few weeks to a few years. Each student independently segmented the identical image and extracted the pixel intensity data. Representative 5x magnification images of each genotype are shown in Figure 4A. Inter-student variation was at or below 4% for all but the lightest-eyed stocks where low pixel intensity values increase variance as a percent of the total to 6% (Figure 4B).

**Figure 4.**
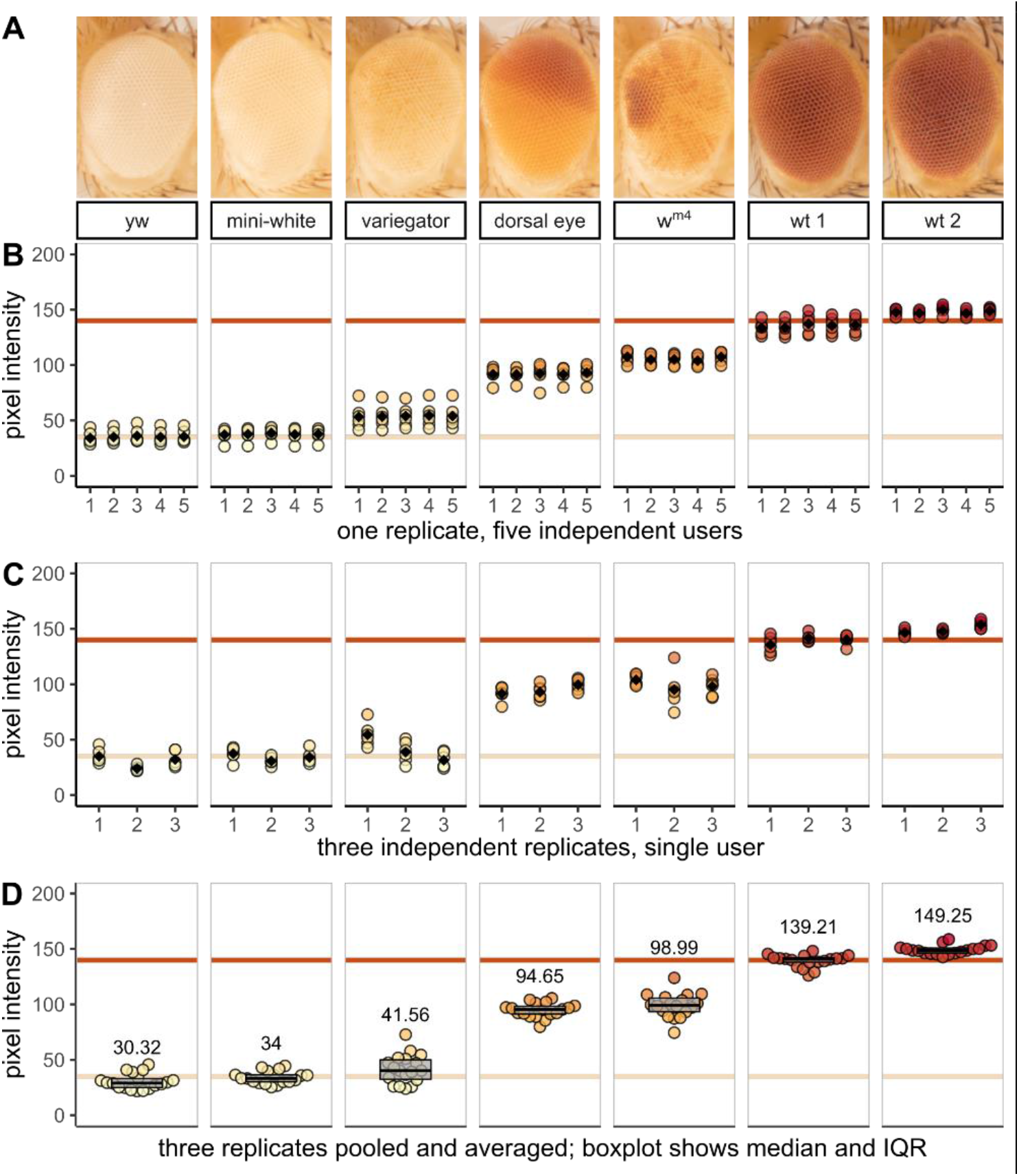
Morphometric analysis of eye pigment data demonstrates precision, accuracy, and reproducibility. Female yw flies were crossed with males from one of seven fly stocks with different eye color phenotypes. Three independent biological replicates were collected for analysis. (**A**) representative 5x magnification images of each stock or genotype in this study; (**B-D**) quantitative data extracted from experimental images. Each filled circle represents the mean pixel intensity of one eye; the beige and red horizontal lines denote the average pixel intensity value for white and wild-type stocks respectively. Black diamonds in (**A**) and (**B**) and numerical values in (**C**) represent the group means. Boxplots in (**C**) show interquartile range with the median as a black horizontal line.

Photographs of three independent biological replicates were analyzed by a single experienced researcher. Biological variation is evident across the three replicates, but even modest inter-group phenotypic differences are detectable (Figure 4C). Flies from all three biological replicates are combined in a bee swarm plot to show the distribution and mean of all data points and a box plot to summarize the median and interquartile range (Figure 4D).

Recognition of the importance of data availability and transparency has grown alongside the widespread accessibility of powerful open-source computational tools. Diverse graphical and statistical approaches can now replace tools like bar graphs and null hypothesis significance testing with detailed data visualizations (Weissgerber et al. 2019) that emphasize effect size as opposed to mere statistical significance (Ho et al. 2019). To this end, our acquisition preserves pixel-level quantitative data for each sample, visualized in ridgeline plots in Figure 2 and dot plots in Figures 3 and 4.

### Other applications and future directions

As demonstrated above, the workflow described here can be used for precise quantitative analysis of eye color phenotypes in *D. melanogaster* and, by extension, for other morphometric data on the shape, size, and color of features as small as 10 µm. For example, studies of pupal size (Sriskanthadevan-Pirahas et al. 2022), wing vein patterning (Alba et al. 2021), photoreceptor neurodegeneration (Dalton et al. 2022), or eye mosaic or clonal analysis (Merkle et al. 2023) could be accelerated by the capacity to image and measure dozens of samples in a single image. Photographic analysis of pigmentation also preserves population variation and individual spatial distribution that could facilitate studies of spatially patterned traits like pigment gene expression (Akiyama et al. 2022). It could also be used for genome-wide association and quantitative trait locus mapping, such as in recent studies of the genetic architecture of abdominal pigmentation across populations of *D. melanogaster* (Dembeck et al. 2015) or the *Drosophila* genus (Ng et al. 2008; Signor et al. 2016). In many of the above examples, sample preparation involves a labor-intensive combination of immersion, dissection, or mounting, which are dramatically simplified here. Additional automation could also be implemented, for example, to program a motorized stage to capture high magnification images of many samples with high reproducibility and low hands-on time.

A variety of computational methods have been developed to automatically segment images of fly eyes or individual ommatidia (Currea et al. 2023), and to reproducibly measure photoreceptor neurodegeneration (Diez-Hermano et al. 2015; Iyer et al. 2016; Diez-Hermano et al. 2020). But using eye morphology as a readout for gene expression and interaction means that in any given experiment, the eyes can have irregular phenotypes, confounding automated image segmentation algorithms that rely on consistent size, shape, color, or position. Where powerful computers with large training datasets struggle, a student can quickly and reliably identify a fly’s eye under the microscope on their first day in the lab, even if that eye is a non-standard shape, size, or color.

While Figure 4A demonstrates that students always identify and segment fly eyes with a high level of precision, this step is labor-intensive and perhaps can soon be automated (Kirillov et al. 2023; Ma and Wang 2023). Focus-stacked macro images like the ones described here could then be fed into computational pipelines like those used to screen pharmaceutical candidate compounds (Stirling et al. 2021).

## Data Availability

Fly stocks are available upon request. All design files, raw data, code, and source images described in this manuscript are available at https://doi.org/10.6084/m9.figshare.25066367 (Arsham 2024).

## Acknowledgments

Stocks obtained from the Bloomington Drosophila Stock Center (RRID:SCR_006457; NIH P40OD018537) were used in this study. We thank the TRiP at Harvard Medical School (NIH/NIGMS R01-GM084947) for creating the transgenic RNAi fly stocks. Funding for this work was provided by Bemidji State University’s New Faculty Scholarship and Innovation and Professional Improvement Grants. We are grateful for the support of the North Hennepin Community College and Bemidji State University communities, and for the ideas and hard work of lab members. Jamie Pipkin was instrumental in designing, prototyping, and fabricating the lighting system and sample tray. We thank Nhi Vuong, Jessica Xiong, Hannah Rhee, and Riley Reed for pilot pigment data analysis. We are indebted to Dr. Sarah C.R. Elgin and her lab whose work is the foundation of the projects described here. This work was conducted on the homelands of Ojibwe and Dakota people to whom the US Government has not fulfilled its legal and financial obligations established in treaties of 1851 and 1863.

